# Heterologous Cas9 and non-homologous end joining as a Potentially Organism-Agnostic Knockout (POAK) system in bacteria

**DOI:** 10.1101/601971

**Authors:** Isaac N. Plant

## Abstract

Making targeted gene deletions is essential for studying organisms, but is difficult in many prokaryotes due to the inefficiency of homologous recombination based methods. Here, I describe an easily modifiable, single-plasmid system that can be used to make rapid, sequence targeted, markerless knockouts in both a Gram-negative and a Gram-positive organism. The system is comprised of targeted DNA cleavage by Cas9 and error-prone repair by Non-Homologous End Joining (NHEJ) proteins. I confirm previous results showing that Cas9 and NHEJ can make knockouts when NHEJ is expressed before Cas9. Then, I show that Cas9 and NHEJ can be used to make knockouts when expressed simultaneously. I term the new method Potentially Organism-Agnostic Knockout (POAK) system and characterize its function in *Escherichia coli* and *Weissella confusa*. First, I develop a novel transformation protocol for *W. confusa*. Next, I show that, as in *E. coli*, POAK can create knockouts in *W. confusa*. Characterization of knockout efficiency across *galK* in both *E. coli* and *W. confusa* showed that while all gRNAs are effective in *E. coli*, only some gRNAs are effective in *W. confusa*, and cut site position within a gene does not determine knockout efficiency for either organism. I examine the sequences of knockouts in both organisms and show that POAK produces similar edits in both *E. coli* and *W. confusa*. Finally, as an example of the importance of being able to make knockouts quickly, I target *W. confusa* sugar metabolism genes to show that two sugar importers are not necessary for metabolism of their respective sugars. Having demonstrated that simultaneous expression of Cas9 and NHEJ is sufficient for making knockouts in two minimally related bacteria, POAK represents a hopeful avenue for making knockouts in other under-utilized bacteria.

## Introduction

Targeted knockouts are the cornerstone of genetics, and the ease of obtaining knockouts can determine how well an organism is studied^1,2^. This has been illustrated in mammalian systems by the messianic reception given to CRISPR/Cas9^3^. However, while it used to be extremely difficult to generate knockouts for many eukaryotic systems, knockout techniques are still limiting in the majority of known bacteria (and prokaryotes more generally)^4^. Despite the fact that CRISPR/Cas9 and host DNA repair can be used to make knockouts in eukaryotic systems without any additional pieces, this is not the case in most bacteria.

Double Stranded Breaks (DSB) in DNA can be repaired by two mechanisms: Homologous Recombination (HR) and Non-Homologous End Joining (NHEJ)^5,6^. HR repairs DSBs using an unbroken template. It duplicates the unbroken template exactly, resulting in error-free repair. HR can be used to generate knockouts or make modifications to the chromosome, but this requires supplying a modified DNA sequence for the cell to use as a template (Fig. **??**). NHEJ, on the other hand, takes two ends of DNA and glues them together. NHEJ systems are generally composed of two proteins, Ku and Ligase D (LigD). Ku binds to free DNA ends as a hexamer and recruits LigD, which ligates two DNA ends together (Fig. 1A). If this occurs immediately after a DSB, the repair may be error-free. However, if there is any damage to the ends of the DNA, such as exonuclease chew back, NHEJ repair will be error prone. As such, a DSB plus NHEJ can result in knockouts. Up until recently, this method was not practical in any organism as it was prohibitively difficult to make targeted DSBs. CRISPR/Cas9 has alleviated the problem of making target DSBs. Unfortunately, unlike eukaryotes, most bacterial species do not have NHEJ^4^.

**Figure 1:**
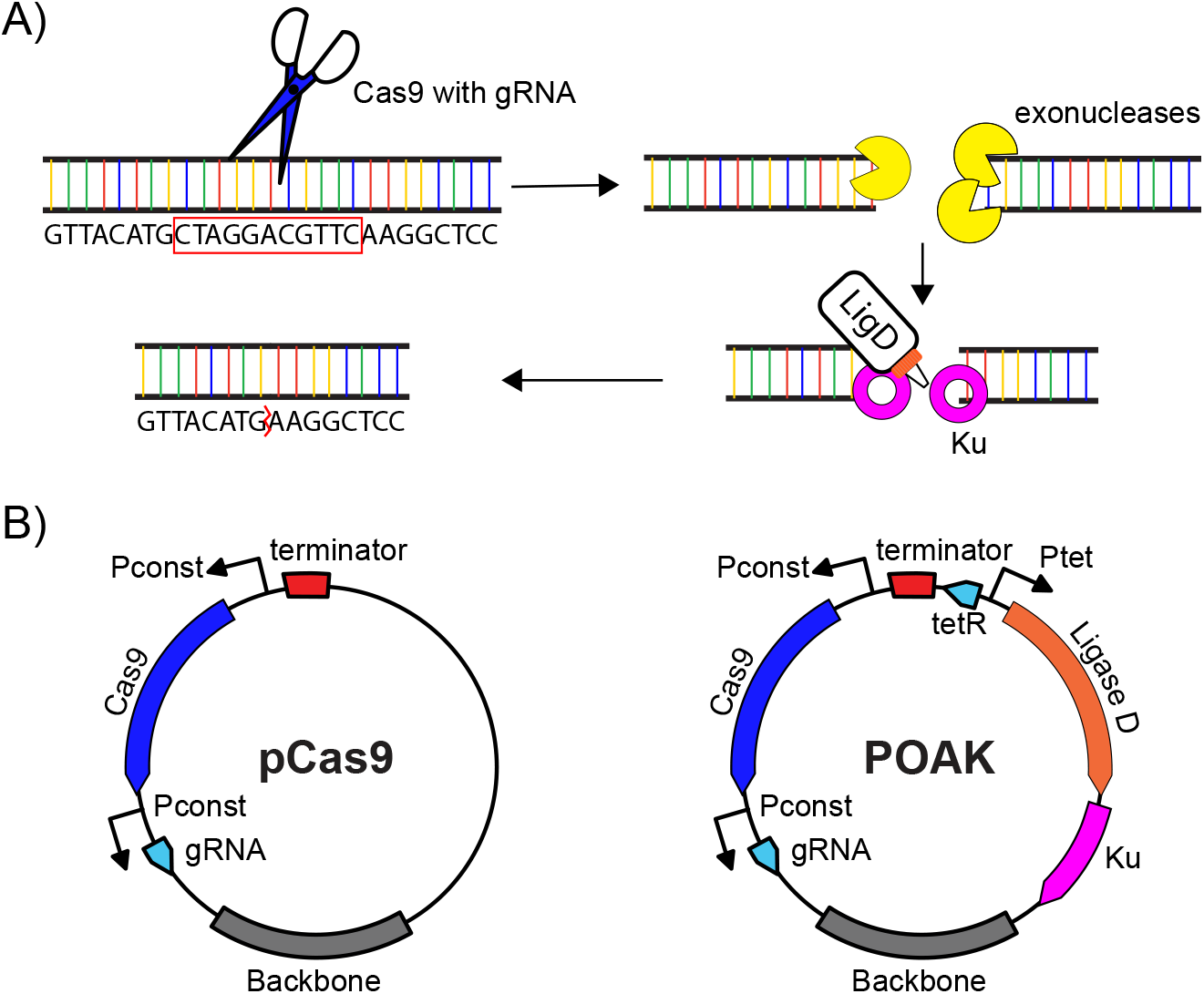
Molecular biology and plasmid maps of pCas9 and POAK. **A)** The molecular biology of NHEJ. A double strand DNA break (DSB) occurs, potentially caused by Cas9 (with gRNA) cutting at a target site. Exonucleases may chew back the ends of the DNA before Ku binds to the ends of the DNA and recruits Ligase D (ligD), which ligates the ends of the DNA. If chew back occurs, the process results in a deletion. **B)** Plasmid maps of pCas9 and POAK.

As a result, HR mediated approaches have been the only approach for targeted knockouts in most bacteria, with other strategies being even more limited (Table 1). In most organisms, however, HR is inefficient and requires long homology arms. In permissive species, a minimum of 500 base pairs may be used, but in many species thousands of base pairs may be necessary^7^. In general, this means making knockouts is very difficult. Certain systems, such as the Lambda Red system in *Escherichia coli*, have greatly decreased the length of homology needed and increased the efficiency of recombination^8^. However, these single strand recombination technologies are derived from phages and are species specific. As a result, they are time consuming to develop and not portable to new organisms.

**Table 1:** Comparison of knockout technologies.

|  | Sequence Specific | Broad Host Range | Efficient | Markerless | Multiple Knockouts |
| --- | --- | --- | --- | --- | --- |
| Mutagenesis Screens<sup>9</sup> |  | ✓ | ✓ | ✓ | ✓ |
| Transposon Mutagenesis<sup>10,11</sup> |  | ✓ | ✓ |  |  |
| Homologous Recombination<sup>7</sup> | ✓ | ✓ |  | - | - |
| Lambda Red<sup>8</sup> | ✓ |  | ✓ | - | - |
| Su *et al.*<sup>12</sup> | ✓ |  | - | ✓ | ✓ |
| POAK | ✓ | ✓ | ✓ | ✓ | ✓ |

A ✓ indicates the technology has the property, a - that it has the property either under specific conditions or only with additional components, and an empty box indicates that the technology does not possess the property.

These difficulties are what have made the use of CRISPR/Cas9 so appealing for use in eukaryotic cells (which have NHEJ). When Cas9 and a gRNA are introduced into the cell, they create a targeted DSB. If repaired through an error free repair mechanism, such as HR, the target sequence can simply be re-cleaved and the cell either dies (if Cas9 cleavage is significantly more efficient than HR) or survives with a wild type sequence (if HR is more efficient). If, instead, the repair mechanism is error prone, such as with NHEJ, the DSB can be repaired with errors that mutate the target sequence and therefore prevent further cleavage (Fig. 1A). These errors are generally deletions (due to exonuclease activity), which makes Cas9 with NHEJ an effective knockout generation system.

For this to work in most bacteria, NHEJ must be provided *in trans*. Significant progress has been made towards making large deletions in *E. coli* by expressing both Cas9 and NHEJ heterologously^12^. That system, however, depends on *E. coli* specific technologies, such as multiple characterized plasmids, that are not available for many bacteria of interest. As such, I sought to interrogate Cas9 paired with NHEJ and determine whether they could be combined into a system that could make knockouts in arbitrary bacteria. Minimally, such a system would have the following features.

1. Both Cas9 and the NHEJ proteins should function in a wide variety of organisms
2. The system should function when combined on a single plasmid
3. The system should be able to create knockouts when expressed simultaneously
4. The system should be easily modifiable to work in any desired organism

Cas9 has been shown to be functional in a wide variety of organisms, both prokaryotic and otherwise^13^. Bacterial NHEJ systems have been less extensively studied^14–16^. Fortunately, NHEJ systems are present in a diverse group of bacteria, from *Mycobacterium tuberculosis* to *Pseudomonas aeruginosa*, and some systems have been used for engineering in their native context^17,18^. Relatively little work has been done to express these systems heterologously, but one study showed NHEJ was able to circularize transformed linear DNA in *E. coli* ^19^ and another *E. coli* study showed NHEJ can repair DSBs caused by Cas9^12^.

In this latter study, Su *et al*. demonstrated that Cas9 and NHEJ could be used in *E*.*coli* to create knockouts, particularly large deletions (Table 1). However, their system depends on NHEJ being expressed in cells prior to expression of the Cas9 and gRNA. In turn, the system also requires the use of multiple plasmids. These features are problematic for use in other bacteria, which often have a minimal number of characterized plasmids^20–22^.

After confirming the dual plasmid results from Su *et al*., I sought to understand whether Cas9 and NHEJ were functional in *E. coli* when expressed from a single plasmid. I also tested Cpf1, a Cas9 like protein that creates a different type of DSB, to see if the type of DSB affects the efficiency of NHEJ. I used a constitutively expressed Cas9, and this had the practical result of also testing whether Cas9 and NHEJ could create knockouts when expressed simultaneously. I then attempted to make knockouts in an organism that had been minimally characterized. For this task, I chose *Weissella confusa*.

*W. confusa* is a Lactic Acid Bacteria (LAB) in the *Leuconostocaceae* family^23,24^. Species in the *Weissella* genus have been studied for biotechnological applications, but these investigations have been limited by the lack of genetic tools. *W. confusa*, in particular, has been transformed, but has had minimal further characterization^25^. I used *W. confusa* as a test case for the organism agnosticism of my single plasmid knockout system.

In both *E. coli* and *W. confusa*, I characterized the effectiveness of POAK, the nature of the knockouts POAK produced, as well as whether the POAK plasmid was stable. As an example of the value of exploring new organisms, I used the knockouts to examine sugar metabolism in *W. confusa*. I hope that POAK will help lower the barrier to making sequence specific, markerless knockouts in arbitrary organisms.

## Results

### Cas9 and NHEJ in *E. coli*

Cas9 and NHEJ were confirmed to be functional in *E. coli*. Due to the limited work that had been done to examine targeted cutting by Cas9 and repair by NHEJ in *E. coli*, I first sought to confirm previous results showing Cas9 could create knockouts in *E. coli* that already expressed the NHEJ proteins. I additionally tested Cpf1. Whereas Cas9 creates blunt end DSBs, Cpf1 creates DSBs with sticky ends, which could conceivably impact the behavior of the NHEJ system. I transformed *E. coli* with either an empty vehicle plasmid (pVeh) or a plasmid encoding constitutively expressed NHEJ (pNHEJ), and subsequently transformed in Cas9 (pCas9sp) or Cpf1 (pCpf1sp) plasmids that contained zero, one, or two gRNAs. Relative survival and knockout efficiency were measured (Fig. 2). In the absence of NHEJ, Cas9 and Cpf1 are highly lethal (Supp. Fig. 9A and B). When NHEJ is present, it significantly increases the survival of *E. coli* transformed with the nucleases plus gRNA (Supp. Fig. 9A and B). To assay for knockout efficiency, gRNAs were targeted to *galK*, a gene essential for galactose metabolism. When plated on MacConkey-galactose agar, colonies with *galk* turn red, while colonies without *galK* turn white. Cas9 or Cpf1 transformed with gRNAs creates a significant percentage of knockouts in an NHEJ strain (Supp. Fig. 9A and B), confirming the work of Su *et al*..

**Figure 2:**
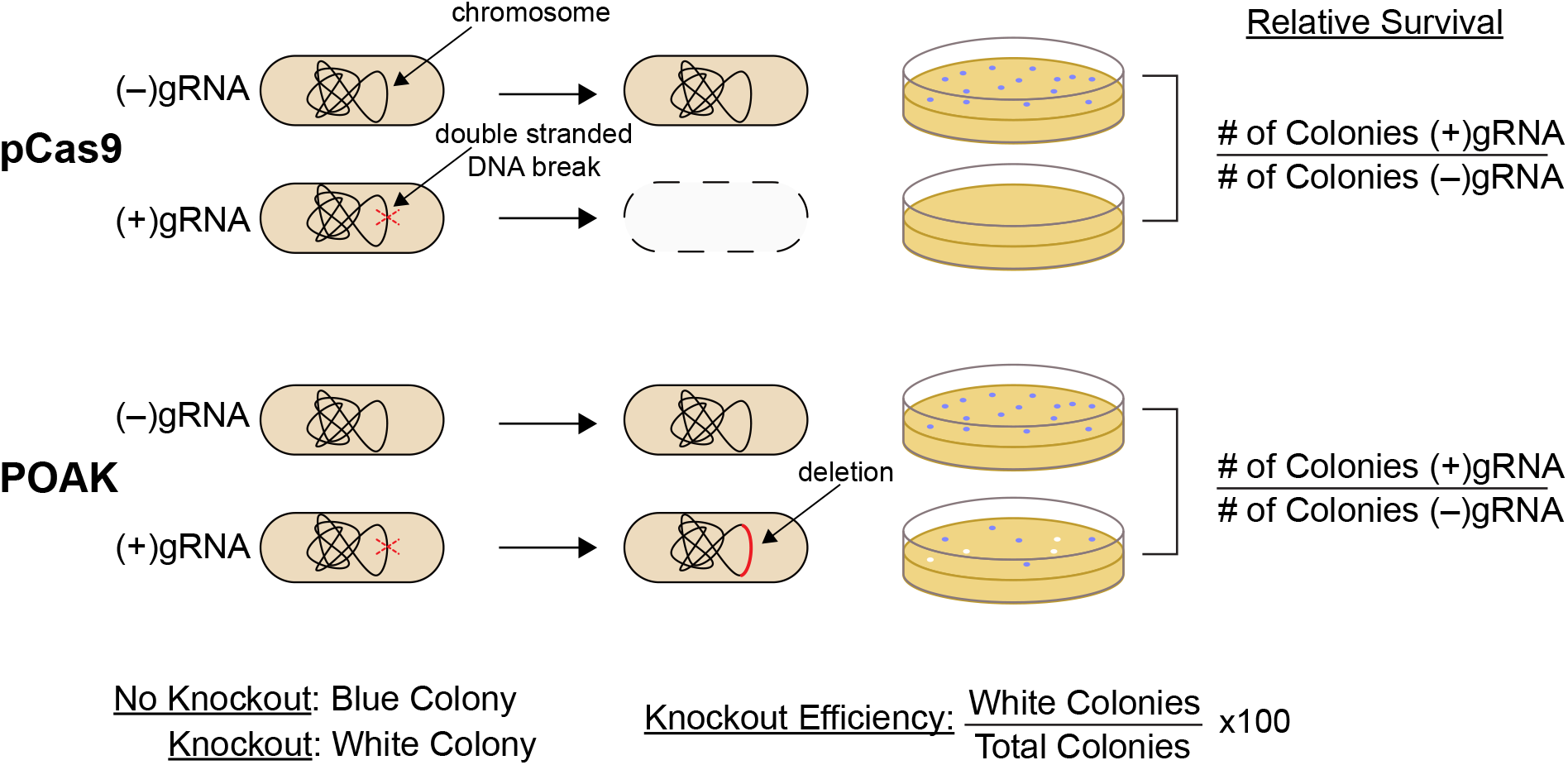
POAK experimental design. Schematic of the experimental design for testing pCas9 and POAK constructs. Transformation efficiency of pCas9/POAK with gRNA is compared to without gRNA to determine relative survival. Colony phenotype, in this illustration blue vs white colonies, is used to determine knockout efficiency.

POAK was made by combining Cas9 and NHEJ on a single plasmid (Fig. 1B). To make Cas9 and NHEJ into a Potentially Organism-Agnostic Knockout (POAK) system, I first combined Cas9, a CRISPR array (for gRNA production), and NHEJ onto a single plasmid. Two versions were created-one for Gram-negative and one for Gram-positive bacteria. To construct the Gram-negative plasmids, a CRISPR array with BsaI sites for gRNA and Cas9 from *Streptococcus pyogenes* (or Cpf1 with its CRISPR array) were combined onto plasmids, and then that plasmids was combined with Gram-negative broad host range origin of replication (Bbr1)^26^ and kanamycin resistance (KnR) to form pCas9 and pCpf1. NHEJ was then placed into the plasmid under tetR control (from the Tn10 transposon)^27^ using two codon optimized gBlocks to create gnPOAK and gnPOAK_Cpf1 (Fig. 1B). To create plasmids for use in Gram-positive organisms, the Bbr1 and KnR on pCas9 and gnPOAK were exchanged for a backbone that contained a broad host range Gram-positive origin of replication^28^, the colE1 origin of replication, and erythromycin resistance^29^, to create the intermediate plasmids pCas9temp and POAKtemp. Finally, the promoter for Cas9 in pCas9temp and POAKtemp was replaced with the 200 bp upstream of the *W. confusa* enolase gene (P_wc-eno_), resulting in plasmids pWcCas9 and gpPOAK. These final two plasmids, as well pCas9, pCpf1, gnPOAK and gnPOAK_Cpf1, were used in further experiments. Table 2 lists these plasmid backbones as well as what they are used for in this work. gRNAs were added to these plasmids as needed, and are referred to by the gene name they target followed by a letter representing the organism as well as a number (e.g. chbCe1, xylAw1). gRNA names are not italicized.

**Table 2:** Plasmid backbones.

| Plasmid | Use | Description |
| --- | --- | --- |
| pCas9 | Cas9 vector with gRNA cloning site for use in *E. coli* | PproC(Cas9), pBBR1 origin, KnR |
| gnPOAK | POAK vector with gRNA cloning site for use in *E. coli* | PproC(Cas9), Pteta(ligd, ku), pBBR1 origin, KnR |
| pCpf1 | Cpf1 vector with gRNA cloning site for use in *E. coli* | PproC(Cpf1), pBBR1 origin, KnR |
| gnPOAK_Cpf1 | POAK_Cpf1 vector with gRNA cloning site for use in *E. coli* | PproC(Cpf1), Pteta(ligd, ku), pBBR1 origin, KnR |
| pCas9temp | Precursor to pWcCas9 | PproC(Cas9), pBAV1K backbone |
| gpPOAKtemp | Precursor to gpPOAK | PproC(Cas9), Pteta(ligd, ku), pBAV1K backbone |
| pWcCas9 | Cas9 vector with gRNA cloning site for use in *W. confusa* | Pwc-eno(Cas9), pBAV1K backbone |
| gpPOAK | POAK vector with gRNA cloning site for use in *W. confusa* | Pwc-eno(Cas9), Pteta(ligd, ku), pBAV1K backbone |

The plasmid backbones used in the main text of the study. gRNAs are appended when they are added to the backbone. So, the pWcCas9 backbone with the galKw1 gRNA would be pWcCas9galKw1. Full lists of the plasmids, strains, and gRNAs used in this study can be found in Supplemental Material and Methods.

NHEJ expressed simultaneously with Cas9 rescues *E. coli* and results in knockouts. To test whether NHEJ expressed at the same time as Cas9 was sufficient to reduce Cas9 lethality and create knockouts, I transformed gnPOAK and gnPOAK_Cpf1 into *E. coli* either with or without a gRNA targeting *galk*. NHEJ is capable of rescuing *E. coli* from Cas9 targeting, but not from Cpf1 targeting (Fig. 3A). Likewise, gnPOAK creates significantly more knockouts than pCas9, while gnPOAK_Cpf1 does not create significantly more knockouts than pCpf1 (Fig. 3B). The relative survival and knockouts efficiency are significantly lower for the gnPOAK system than the dual plasmid system. Further, there is high variability in both survival as well as knockout efficiency in the single plasmid system.

**Figure 3:**
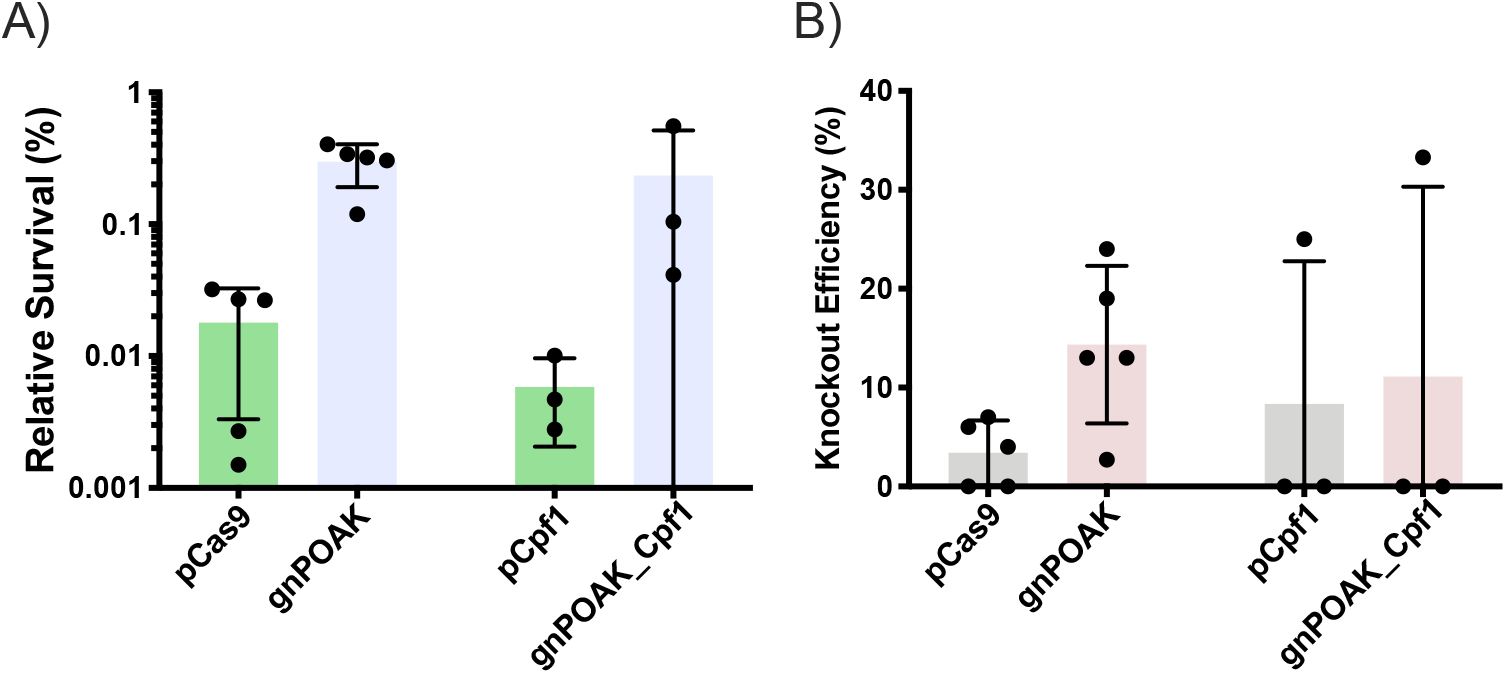
NHEJ increases survival and knockouts from DSBs caused by Cas9 but not Cpf1 in *E. coli*. pCas9, pCpf1, gnPOAK, and gnPOAK_Cpf1 were transformed into *E. coli* with galKe9 (for pCas9 and gnPOAK) or galKe1F (for pCpf1 and gnPOAK_Cpf1). **A)** Relative survival of transformations of pCas9galKe9 and gnPOAKgalKe9 (n=6) and of pCpf1galKe1F and gnPOAK_Cpf1galKe1F (n=3). Bars are the mean, and errors bars represent the standard deviation. **B)** Knockout efficiency of transformations of pCas9galKe9 and gnPOAKgalKe9 (n=6) and of pCpf1galKe1F and gnPOAK_Cpf1galKe1F (n=3). Bars are the mean, and errors bars represent the standard deviation.

### Cas9 and NHEJ in *W. confusa*

I found that *W confusa* can be transformed with high efficiency using a simple electroporation protocol. The only previously published transformation protocol for *W. confusa* involves making protoplasts through enzymatic digestion of the cell wall^25^. Here I developed a *W. confusa* electrocompetent preparation as a room temperature (RT) protocol, although certain steps showed sensitivity to overheating (Fig. 4A). During the wash steps, if cells were spun down in a non-temperature-controlled centrifuge that was being heavily used, transformations routinely failed (data not shown). I suspect this is due to overheating, and in these cases the cell pellets were looser and significant debris remained in the media, possibly indicating cell lysis. This is supported by *W. confusa’s* rapid death upon heat shock (Supp. Fig. 10). The electroporation itself was also sensitive to overheating. Cells electroporated in ice cold (0°C) cuvettes showed significantly higher transformation rates than cells electroporated in RT cuvettes (Fig. 4B). Samples electroporated in RT cuvettes had released significant amounts of genomic DNA (which could be observed by eye during the rescue step of the transformations), suggesting lysis occurred during electroporation.

**Figure 4:**
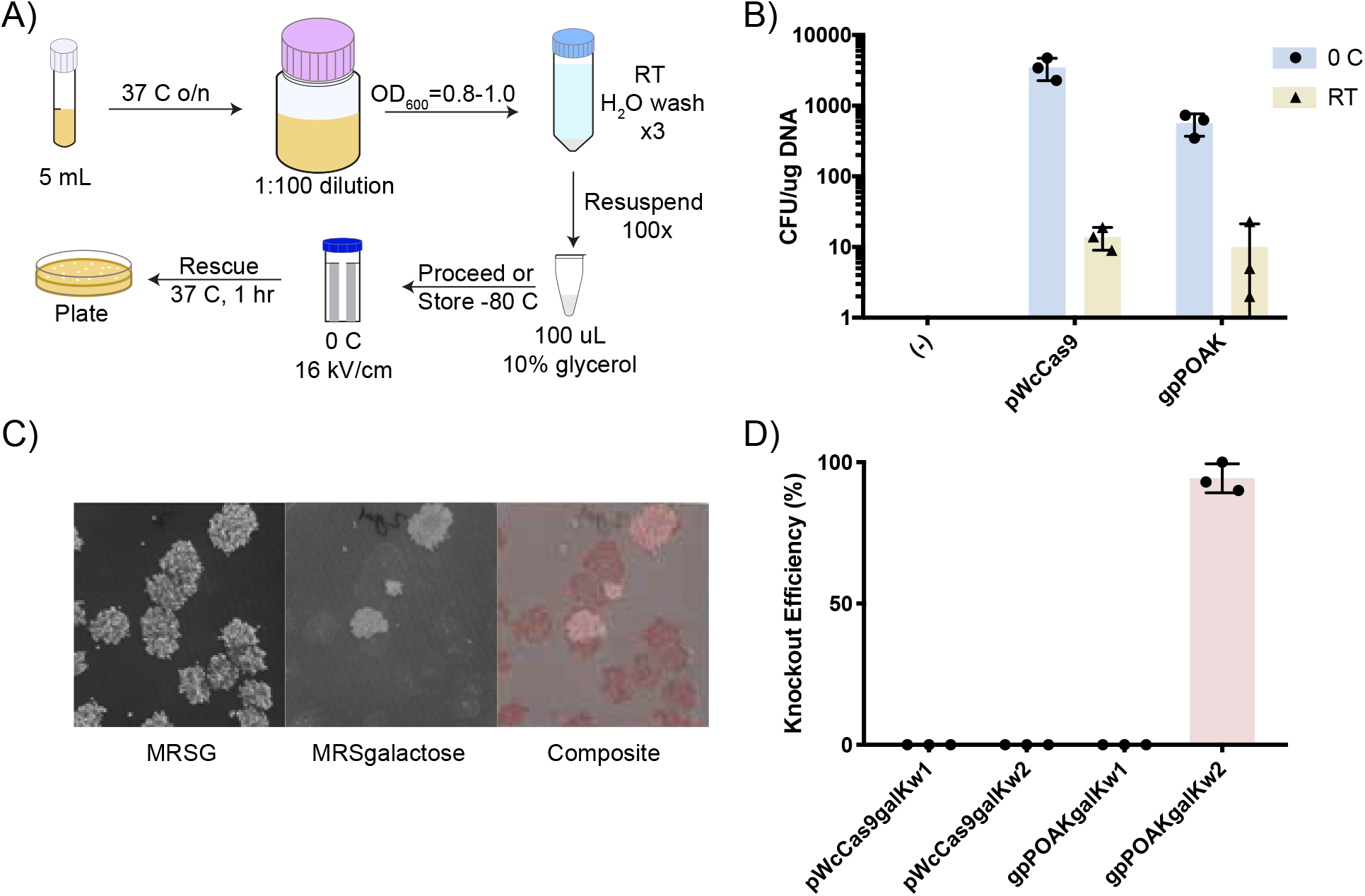
Transformation and knockouts using POAK in *W. confusa*. **A)** Schematic of transformation protocol for *W. confusa*. An overnight culture is back-diluted 1:100 and grown until OD_600_ 0.8-1. The culture is washed three times with RT water, resuspended at 100x concentration, then electroporated at 16 kV*/*cm. **B)** Transformation efficiency of electroporation in *W. confusa* with two different plasmids, pWcCas9 and gpPOAK, and at two different cuvette temperatures, 0°C and RT. Bars show the mean of three transformations and error bars represent the standard deviation. **C)** Knockouts after transformation of *W. confusa* with gpPOAKgalKw2. Replica plating of a representative transformation onto MRSG, MRSgalactose, and a composite of the two is shown. **D)** Knockout efficiency after transformation with pWcCas9galkw1, gpPOAKgalkw2, pWcas9galkw2, and gpPOAK-galkw2. Bars are the mean of three transformations and error bars show standard deviation.

POAK can make knockouts in *W. confusa*. To test whether gpPOAK could make knockouts in *W. confusa*, I designed gRNAs targeted either 100 bp (galKw1) or 200 bp (galKw2) downstream of the start codon in the *W. confusa galK* gene. A knockout in *galK* should result in a loss of galactose metabolism. To determine if transformants could metabolize galactose, I developed a replica plating technique for *W. confusa*. In short, this involved replica plating transformation plates onto a second set of selective plates (to mimic restreaking of colonies) then replica plating the second set of selective plates onto plates selective for galactose metabolism (MRS media with galactose, MRSgalactose) and non-selective for galactose metabolism (MRS media with glucose, MRSG). This allows for assaying of sugar metabolism knockouts (Fig. 4C). gpPOAK’s ability to make knockouts is gRNA dependent and NHEJ dependent in *W. confusa* (Fig. 4D). To test this, I transformed pWc-Cas9galKw1, pWcCas9galKw2, gpPOAKgalKw1, and gpPOAKgalKw2 into *W. confusa*. Of these, only gpPOAKgalKw2 created observable knockouts, suggesting that both NHEJ and specific gRNAs are necessary.

pWcCas9 does not efficiently cut the *W. confusa* genome. To test the efficiency of Cas9 in *W. confusa*, the relative survival of pWcCas9 with galKw1 or galKw2 was assayed (Supp. Fig. 11). Despite leading to knockouts when used with gpPOAK, neither gRNA decreased the survival of *W. confusa* below 10% whether used in pWcCas9 or gpPOAK. In contrast, Cas9 with gRNA efficiently reduces survival in *E. coli* (Fig. 3A).

### Comparison of POAK Behavior in *E. coli* and *W. confusa*

POAK knockouts can be made by targeting any part of a gene in *E. coli* and *W. confusa*. To test the effect of cut location on knockout efficiency, I transformed *E. coli* and *W. confusa* with POAK plasmids that targeted regions approximately 100 bp apart along each organism’s respective *galK* sequence. The percentage of colonies that had a Δ*galK* phenotype was assayed by MacConkey-Galactose plates or replica plating on MRSgalactose for *E. coli* and *W. confusa* respectively. Knockouts occurred for gRNAs targeted across the gene in both organisms, although all *E. coli* gRNAs produced knockouts (Fig. 5A) while only ∼50% of *W. confusa* gRNAs produced knockouts (Fig. 5B). For gRNAs that produced knockouts, there is significant variability in the knockout efficiency, ranging from 10% to 40% in *E. coli* and from 4% to 95% in *W. confusa*. The highest knockout efficiencies do not cluster around known GalK actives sites.

**Figure 5:**
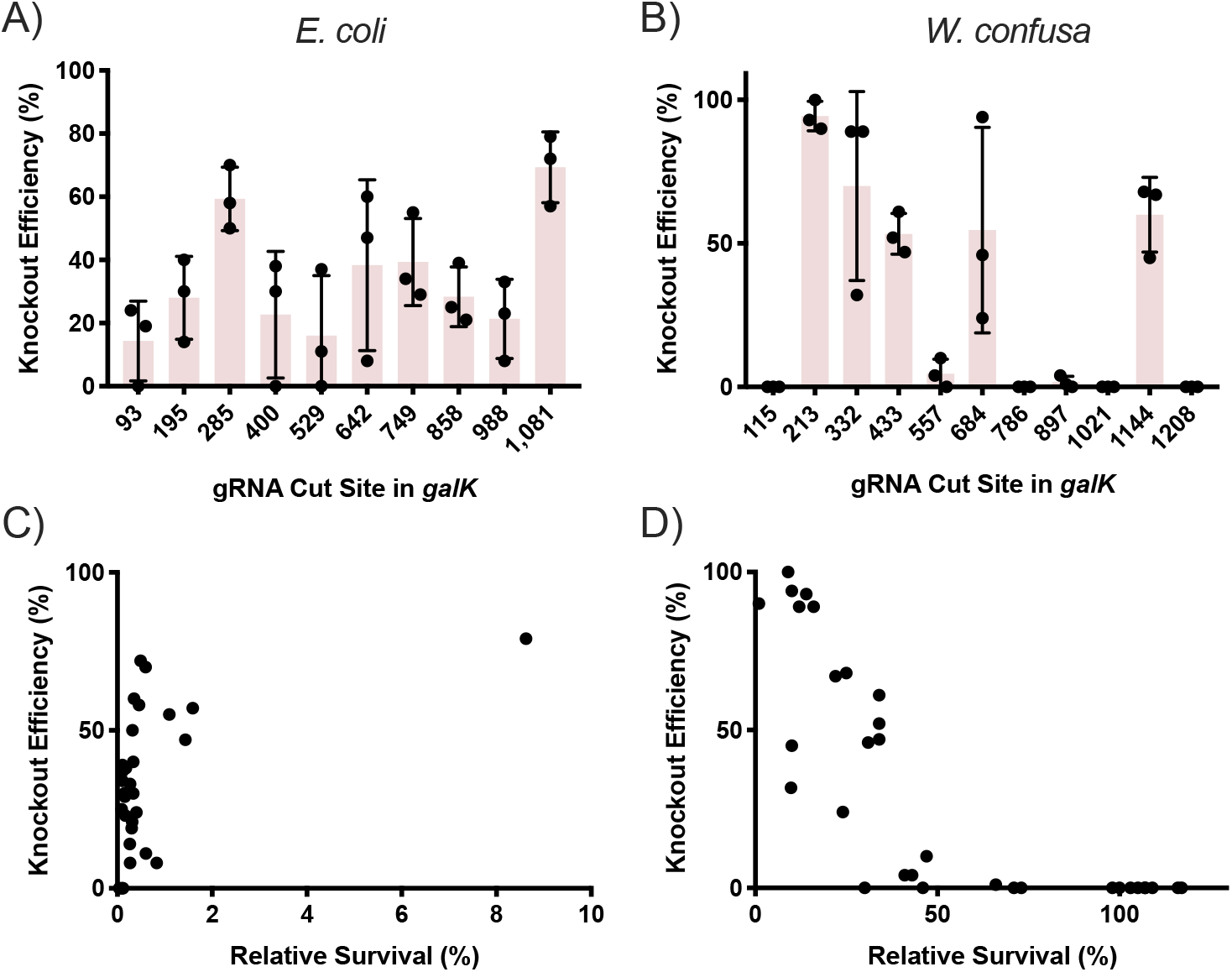
Effect of cut position and relative survival on POAK knockouts in *E. coli* and *W. confusa*. **A)** Knockout efficiency observed on MacConkey+galactose plates when *E. coli* is transformed with gnPOAK containing a gRNA that cuts at the indicated position of *galK*. Bars are the mean of three transformations, error bars represent the standard deviation. **B)** Knockout efficiency observed by replica plating when *W. confusa* is transformed with gpPOAK containing a gRNA that cuts at the indicated position of *galK*. Cut sites 115 bp and 213 bp are gRNAs galKw1 and galKw2 respectively. Bars are the mean of three transformations, error bars represent the standard deviation. **C)** Knockout efficiency vs. relative survival of all transformation in **A. D)** Knockout efficiency vs. relative survival of all transformation in **B**.

POAK knockout efficiency is correlated to Cas9 efficiency in *W. confusa* but not in *E. coli*. To determine why all of the *E. coli* gRNAs produced knockouts but not all of the *W. confusa* gRNAs did, I compared the relative survival to knockout efficiency for all transformations. The relative survival of *E. coli* when transformed with gnPOAK bares no relationship to the number of knockouts that will be produced (Fig. 5C). *W. confusa*, however, has a clear relationship between relative survival and knockout efficiency, with no transformation that had above 50% survival producing knockouts. Survival rates were not correlated with the location of the cut site within the gene for either *E. coli* or *W. confusa*, but individual gRNAs do show similar levels of relative survival across the three transformations (Supp. Fig. 12).

To understand the type of knockouts that POAK creates I targeted five different sugar metabolism genes from the *E. coli* and *W. confusa* genomes (Table 3). Briefly, all surviving colonies from each transformation were pooled and genomic DNA collected. The 4 kbp region surrounding the gRNA cut site was amplified and sequenced using an NGS pipeline. Sequence deletions were observed for all targeted *E. coli* genes. In *W. confusa*, sequence deletions were observed in *celbPTSIIC, manPTSIIC*, and *xylA*. No deletions were observed in *maltP*, and *galK* deletions were observed only from the galKw2 gRNA, and not from galKw1.

**Table 3:** Sugar metabolism genes targeted for NGS experiment.

|  | Gene | Expected Function | gRNA |
| --- | --- | --- | --- |
| *E. coli* |  |  |  |
|  | *chbC* | cellobios importer | chbCe1 |
|  | *galk* | galactose kinase | galKe1 |
|  | *lacZ* | beta-galactosidase (lactose metabolism) | lacZe1 |
|  | *manY* | mannose importer | manYe1 |
|  | *xylA* | xylose isomerase | xylAe1 |
| *W. confusa* |  |  |  |
|  | *celbPTSIIC1* | cellobios importer | celbPTSIICw1 |
|  | *galk* | galactose kinase | galKw1 |
|  |  |  | galKw2 |
|  | *maltP* | maltose phosphorylase | lacZw1 |
|  | *manPTSIIC* | mannose importer | manPTSIICw1 |
|  | *xylA* | xylose isomerase | xylAw1 |

POAK produces a range of deletions in *E. coli* and *W. confusa*. Fig. 6 shows all observed deletions from three replicates of *galK* and *xylA* genes in *E. coli* and *galK* and *manPTSIIC* in *W. confusa*. Each targeted gene shows a range of deletions from as small as 7 bp to up to 2500 bp (sample preparation may have precluded observation of significantly larger deletions). Most deletions occur bi-directionally around the cut site. However, in some cases, such as gpPOAKmanPTSIICw1 (Fig. 6D), observed deletions do occur primarily in one direction. This was also observed in gpPOAKxylA1 for individual replicates (Supp. File A.4). There are more observed deletions for *E. coli*, which is consistent with the larger number of recovered transformants. In *W. confusa*, deletions were observed in *galK* from the gRNA galKw2, but not from galKw1, which is consistent with the knockout data (Fig. 4B).

**Figure 6:**
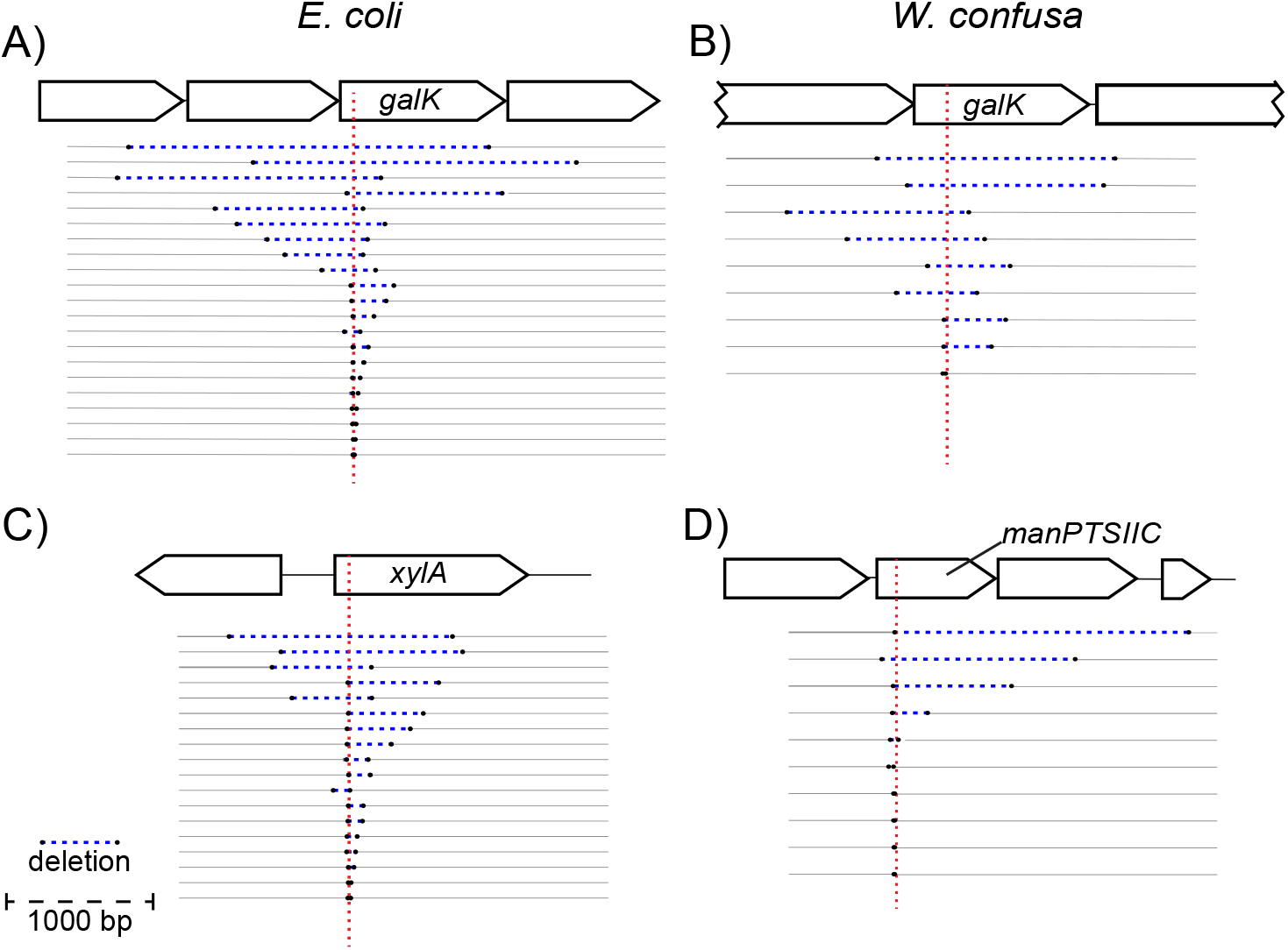
Representative editing by gnPOAK in *E. coli* and of gpPOAK in *W. confusa*. Each horizontal line represents an observed deletion supported by more than five reads. All deletions from three different transformations are shown in each panel. Deletions are indicated by black dots connected by a dotted blue lines, and the Cas9-gRNA cut site is represented by a dotted red line. 1000 bp shown for scale. **A)** Transformations of gnPOAKgalKe1 in *E. coli*. **B)** Transformations of gpPOAKgalKw2 into *W. confusa* **C)** Transformations of gnPOAKxylAe1 in *E. coli* **D)** Transformations of gpPOAKmanPTSI-ICw1 in *W. confusa*

Knockouts of the putative cellobios and mannose importers do not prevent cellobios or mannose metabolism in *W. confusa*. In *E. coli* all gRNAs created sequence deletions and also resulted in loss of sugar metabolism (except for *chbC*, which could not be assayed) (Supp. Fig. 13B). In *W. confusa*, of the four gRNAs that produced observable sequence deletions, only galKw2 resulted in colonies that were completely deficient for metabolism of the relevant sugar (Fig. 4C). xylAwI produced segmented colonies at low rates (Fig. 7B), which is consistent with the gRNA’s high rates of survival (Fig. 7A). To determine whether or not the deletions observed through NGS in *celBPTSIIC* and *manPTSIIC* were preventing *W. confusa* from metabolizing those sugars, knockouts for both genes were reisolated. Knockouts were verified by colony PCR and Sanger sequencing (Supp. Fig. 14). The knockouts both grew when their respective sugar was the sole carbon source (Fig. 7C).

**Figure 7:**
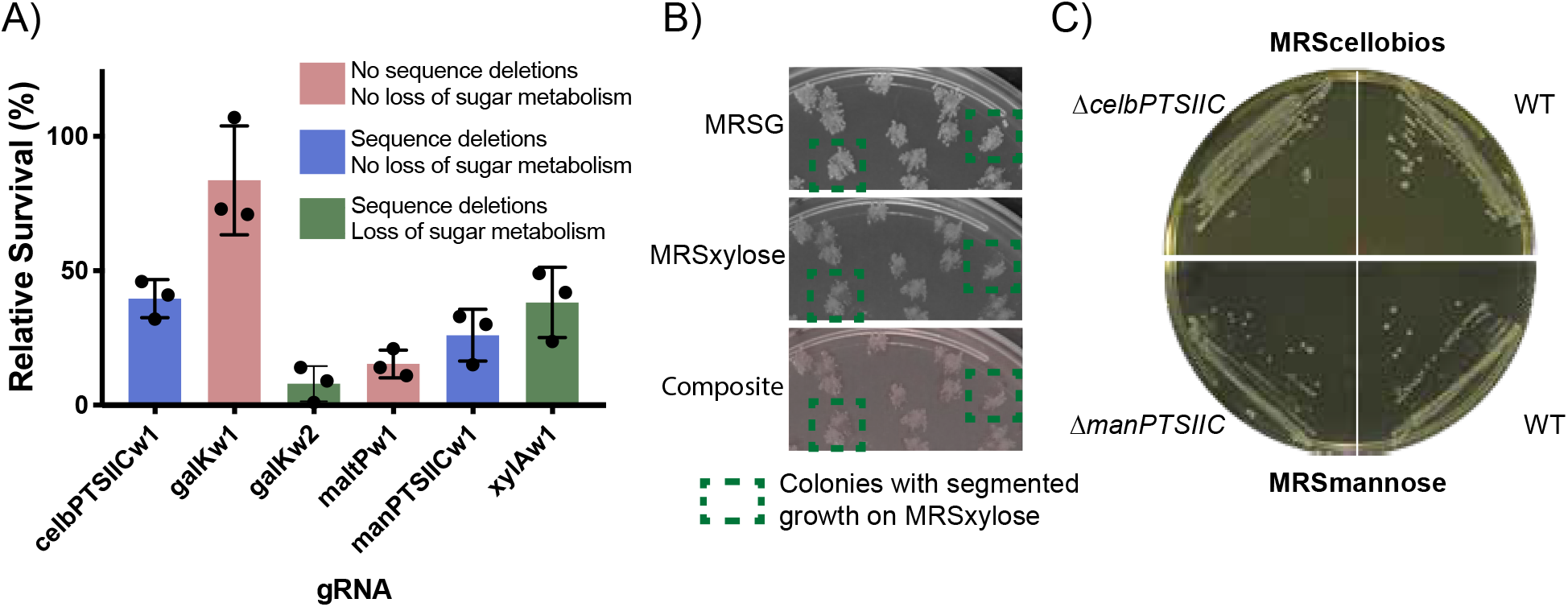
Two genetics knockouts from the *W. confusa* NGS screen do not result in loss of sugar metabolism. **A)** Relative survival for five targeted *W. confusa* genes. Blue bars indicate genes for which deletion mutants were observed in sequencing, but no loss of sugar metabolism was observed after replica plating. Green bars are for genes that showed both sequence deletions and loss of sugar metabolism. Red bars indicate that neither sequence deletions nor loss of sugar metabolism was observed. Bars are the mean of three transformations, error bars represent the standard deviation. **B)** Replica plating of a xylAw1 transformation. Colonies that showed segmented xylose-dependent survival are highlighted. **C)** Knockouts in *celBPTSIIC* and *manPTSIIC* genes grow on plates that require cellobios metabolism (MRScellobios) and mannose metabolism (MRSmannose), respectively. WT *W. confusa* shown on each type of plate.

POAK plasmids rapidly cure from both *E. coli* and *W. confusa*. The components of NHEJ can be toxic to *E. coli* when over-expressed. This appears to due to the effects of Ku, which cause an induction dependent growth defect (Supp. Fig. 15B and D), whereas induction of LigD shows much more subtle effects (Supp. Fig. 15A and C). To test whether this would lead to curing of the plasmid, POAK and Cas9 plasmids were grown in *E. coli* and *W. confusa* in the presence of aTc (which induces the NHEJ proteins) but without selection. Plasmids with NHEJ were cured from over 99% of *E*.*coli* cells and greater than 90% of *W. confusa* cells (Supp. Fig. 16). In *E. coli*, the Cas9 only plasmid was stable during overnight growth, while in *W. confusa* the Cas9 plasmid was lost almost as rapidly as the POAK plasmid. This suggests that while in *E. coli* plasmid instability is caused by NHEJ proteins, in *W. confusa* the plasmid instability is due to the plasmid backbone and not the NHEJ components.

## Discussion

In the introduction, I laid out four conditions that are required for a knockout system to work in arbitrary bacteria. Of these, the outstanding questions were whether Cas9 and NHEJ could create knockouts when expressed simultaneously, and how difficult it would be to modify a Cas9 and NHEJ expression system to work in a chosen organism. In this work, I have shown that Cas9 and NHEJ can create knockouts in both *E. coli* and *W. confusa* when expressed simultaneously. Further, only minimal modifications were required to get the system working in *W. confusa*.

Replacing the promoter for Cas9 was the only species-specific change required for POAK to work in *W. confusa*. To create gpPOAK, the backbone of gnPOAK was changed to an origin of replication (from pBavIK) and antibiotic cassette known to work in Gram-positive baceria. The backbones of both POAK vectors are considered “broad host range”, so these should only require occasional adjustment. The only species-specific adjustment I made to gpPOAK was to add P_wc-eno_ (the 200 bp upstream of the *W. confusa* enolase gene) in front of Cas9. The choice of the enolase promoter was based on data suggesting enolases are often highly expressed in bacteria, not on *W. confusa* specific data. This change was sufficient to produce levels of Cas9 capable of making knockouts. However, it is worth noting that not all gRNAs work in *W. confusa*, while all gRNAs work in *E. coli*. Further, Cas9 and POAK with gRNAs both show much lower rates of survival in *E. coli* than in *W. confusa*. This is not surprising, considering that both the gRNA promoters and the Cas9 promoter used in pCas9 and gnPOAK were optimized for *E. coli* ^30^ while neither the gRNA promoters nor the Cas9 promoter in pWcCas9 or gpPOAK were particularly optimized for use in *W. confusa*. As such, it is likely that further optimization would increase the effectiveness of pWcCas9 and gpPOAK in *W. confusa*.

*W. confusa* is useful for biotechnological and laboratory use. *W. confusa* grows quickly, with a doubling time of 40 minutes at 37°C (Supp. Fig. 17), in both aerobic and anaerobic conditions. Unlike many other LAB and Gram-positive bacteria, *W. confusa* does not lyse in stationary phase. I show it can be transformed with a simple electroporation protocol and is robust to freezing and storage at −80°C. Transformation efficiencies are high enough that I was able to use it as a cloning host for a plasmid that was toxic in *E. coli*. Overall, the organisms ease of use rivals that of *E. coli* and warrants investigation as Gram-positive chassis.

POAK can be used to go from design to sequenced markerless knockout in a week. Once POAK has been validated in an organism, knockouts can be made in under a week, with the marginal time for each additional knockout being about 4 days (Fig. 8). First, gRNAs are designed and ordered (Day 0), then cloned into the appropriate POAK vector and transformed into a cloning strain (Day 1). A colony PCR of the gRNA is sent to confirm the sequence of gRNA insertion and colonies are picked for overnight cultures (Day 2). Sequence validated clones are mini-prepped and transformed into the target organisms (Day 3). Surviving colonies are struck out on selective plates (Day 4+, depending on organism’s growth rate). Colonies are sent in for sequencing of the target locus and colonies are grown overnight in aTc (to induce the NHEJ proteins) and without antibiotics (Day 5+). Sequence confirmed colonies are struck out on non-selective plates (Day 6+). Colonies can then be used as parent strains for further knockouts, although if used immediately, colonies should be picked into both selective and non-selective media to confirm loss of plasmid (Day 7+).

**Figure 8:**
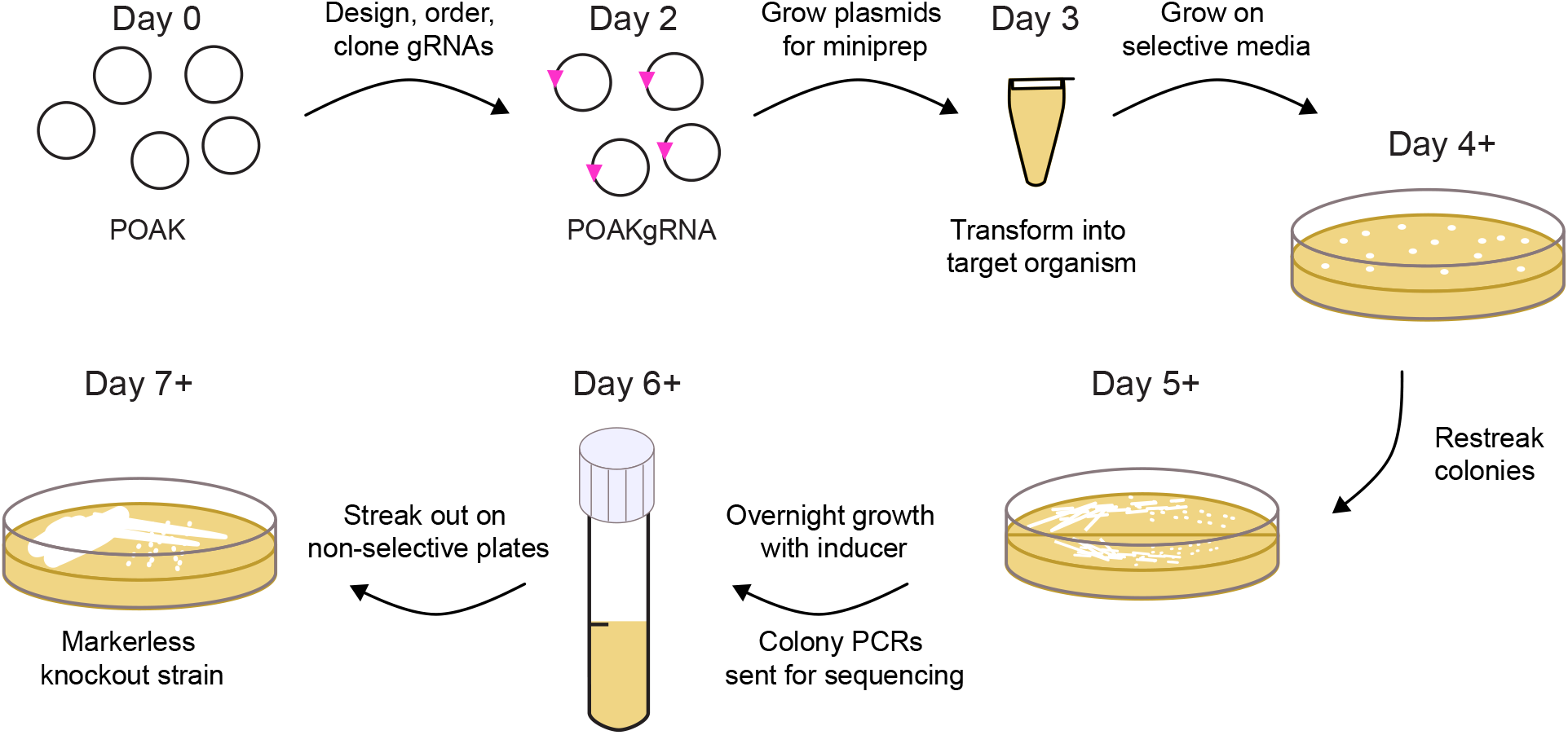
POAKing out genes. From the time gRNAs are designed (**Day 0**) it takes three days until transformation of POAK into desired organism (**Day 3**). After the transformants have grown up (**Day 4+**) it takes three more days before markerless knockouts are obtained (**Day 7+**).

POAK creates a range of deletions. Unlike recombination-based techniques that create a set of uniform transformants, POAK creates a range of mutations. These mutations range from small mutations (SNPs and <10 bp deletions) that can be used for targeted editing, to large mutations (>2000 bp) that can be used to knockout multiple genes at once. A similar range of deletions is observed in both *E. coli* and *W. confusa*. Across multiple replicates, these deletions generally occur symmetrically around the cut site. In certain *W. confusa* replicates, such as xylAw1, there appeared to be asymmetry in the deletion (i.e. most of the deletions primarily include portions either upstream or downstream of the cut site). However, this effect did not occur consistently, and as such I cannot draw any strong conclusions about whether the directionality is a real effect or the conditions under which it might occur.

The deletions created by POAK are markerless, and therefore can be used in applications where downstream repression (e.g. CRISPRi) or the presence of antibiotic cassettes (e.g. HR) are not desirable. This makes POAK ideal for targeting a single gene in an operon, or in organisms for which only a single antibiotic is routinely used. This contrasts with homologous recombination strategies that require additional technologies (such as recombinases) to be rendered markerless. POAK nicely complements CRISPRi, a technique that is based on dCas9 and has been applied in a range of bacteria^31^. Unlike POAK, CRISPRi can create knockdowns of entire operons. The combination of the two techniques has the potential to allow robust genetic interrogation of heretofore recalcitrant organisms.

In this study, I have expanded on previous work that hinted that Cas9 and NHEJ might be capable of creating knockouts in a wide range of bacteria. I showed that simultaneous expression of Cas9 and NHEJ creates knockouts in *E. coli* and in *W. confusa*. The efficiency of making knockouts appears to depend on the efficiency of Cas9 cutting, a feature that can be tuned based on the organism. Regardless of the efficiency of knockout, the resulting sequence edits are similar between both organisms. POAK plasmids are quickly cured in both organisms as well. Taken together, this suggests that I have made progress towards a markerless, sequence specific, broad host range, knockout system. It is my hope that this Potentially Organism Agnostic Knockout (POAK) system will be taken by the community and further developed into a true Prokaryotic Organism Agnostic Knockout (POAK) system.

## Materials and Methods

### Strains and Growth Conditions

Complete strain information can be found in Supplemental Table 4. DH10*β* was used as the default cloning strain, and was routinely grown in Lysogeny Broth (LB) Miller and on LB agar plates. For the Gram-positive shuttle plasmid, DH10*β* was grown in 2xYT broth and on Blood Heart Infusion (BHI) agar. Single gene knockouts from the Keio^32^ collection with the antibiotic resistance removed were used for cloning of plasmids that contained gRNAs targeted to the *E. coli* genome. Experiments were performed in MG1655 (*E. coli K12*) or in DSM 20196 (*W. confusa*). MG1655 was routinely grown in LB and on LB plates. *W. confusa* was grown in Man, Rogosa and Sharpe (MRS)^33^ media with glucose (MRSG) broth and on MRSG plates. All MRS plates were prepared by autoclaving the media *without* sugar, and supplementing filter sterilized sugar to two percent final concentration afterwards.

For *E. coli* galactose, mannose, and xylose metabolism experiments, MacConkey^34^ agar was used with the appropriate sugar supplemented to one percent. *E. coli* lactose metabolism was assayed using LB plates containing X-gal. MRS plates with the appropriate sugar substituted for glucose were used as selective plates for *W. confusa* sugar metabolism experiments. Initially MRS without yeast extract was used, so as to eliminate residual mannose, but the presence or absence of yeast extract was observed to be inconsequential, and so it was included in the MRS used for later replicates. For selective growth, spectinomycin at 100 µg*/*mL, kanamycin at 45 µg*/*mL, and erythromycin at 150 µg*/*mL were supplemented for *E. coli*, and erythromycin at 10 µg*/*mL was supplemented for *W. confusa*.

### Plasmid Construction

A complete list of plasmids can be found in Supplmental Table 5 and plasmid maps can be found in Supplemental File A.1. Golden Gate Assembly^35^ was used for cloning plasmids and inserting gRNAs. For all cloning besides insertion of gRNAs, Q5 Hot Start polymerase was used for amplification of assembly pieces. DNA for the coding sequences of LigD and Ku were ordered as gBlocks from IDT. For Golden Gate reactions, 10x T4 Ligase Buffer (Promega), T4 Ligase (2,000,000 units/mL, NEB), and BSA (10 mg/mL, NEB) were used in all reactions. The appropriate restriction enzyme, either Eco31I, Esp3I, or SapI (Thermo FastDigest), was added. gRNAs were added to plasmids by Golden Gate after annealing and phosphorylating pairs of oligos. For detailed information, see Supplemental Materials and Methods. Briefly, complementary oligos were incubated with T4 Ligase Buffer (NEB) and T4 Poly Nucleotide Kinase (NEB), heated to boiling, and then slowly cooled to room temperature. Annealed oligos were then added to plasmids using Eco31I.

Golden Gate reactions were desalinated using drop dialysis (for a minimum of 10 minutes) and electroporated in DH10*β* Electrocompetent Cells (Thermo Fischer).

### *E. coli* Electrocompetent Preparation and Transformations

*E. coli* was made electrocompetent using a modified standard protocol^36^. *E. coli* was grown in either LB or 2xYT until OD_600_ of between 0.4 and 0.6. Cells were spun down at 4000xG for 10 minutes at room temperature. Cells were then washed twice with 0.5x volume room temperature ddH_2_O and then once with 0.1x volume room temperature ddH_2_O. Cells were re-suspended in approximately 0.001x volume of room temperature ddH_2_O (if competent cells were to be used immediately) or 10% room temperature glycerol (if cells were to be frozen and stored at −80°C). 25 µl of cells were transformed with 2.5 µl of 20 ng*/*µl of mini-prepped plasmid. Electroporations were performed using 0.1 cm cuvettes at 1.8 kV (Ec1) in a BioRad MicroPulser™. Cells were rescued in 972.5 µl SOC supplemented with 200 nm aTc for 1 hour at 37°C. Cells were then plated on agar plates with appropriate selection.

Occasionally, cells prepared using this method are too concentrated and arc, so a no-DNA control electroporation was always performed. If the pulse time was less than 5 ms, cells were diluted until an appropriate pulse time was achieved.

Agar plates were imaged using a macroscope. Colonies were counted manually, and metabolism of the relevant sugar was indicated by red colored colonies on MacConkey plates or blue colonies on LB+X-gal plates. White colored colonies on either media was indicative of no metabolism of the specific sugar.

Replicates are of at least three separate competent cell preparations, except for Supp. Fig. 9 which is three transformations from the same electrocompetent preparation.

### *W. Confusa* Electrocompetent Preparation and Transformations

*W. confusa* was made electrocompetent using a modified version of the above protocol. *W. confsua* was grown, either from a colony or from a 1:100 dilution of an overnight culture, in MRSG until OD_600_ of between 0.8 and 1.0 (often closer to 0.8, due to it being faster). Cells were spun down at 4000xG for 12 minutes at 22°C.^*†*^ Cells were then washed twice with 0.5x volume room temperature ddH_2_O and then once with 0.1x volume room temperature ddH_2_O. Cells were re-suspended in 0.01x volume of room temperature ddH_2_O (if competent cells were to be used immediately) or 10% room temperature glycerol (if cells were to be frozen and stored at −80°C). 100 µl of cells were transformed with 10 µL of 10 ng*/*µl of mini-prepped plasmid. Electroporations were performed using 0.2 cm ice cold cuvettes at 2.5 kV (Ec2) in a BioRad MicroPulser™. Cells were rescued using 900 µl MRSG supplemented with 200 nm aTc for 1.5 hours at 37°C. Cells were then plated on MRSG agar plates with erythromycin and grown at 37°C for two to three days (or until colonies had grown to an adequate size). It is important to use MRS media that has the sugar added after autoclaving. Plates that have been made from MRS autoclaved with sugar significantly reduce transformation efficiency.

Agar plates were imaged using a macroscope. Colonies were counted using the Cell Colony Edge FIJI macro^37^ that had been modified. Results were manually checked for accuracy. Macro is provided as Supp. File A.2. Replicates are of at least three separate competent cell preparations.

Frozen competent cells remain viable for at least nine months (and likely much longer). To use, thaw on ice and proceed as described above.

### *W. confusa* Replica Plating

*W. confusa* was replica plated using either a sterile felt or two to three paper towels from the inside of an unopened stack. Colonies were first replica plated onto MRSG+erythromycin and then grown at 37°C overnight. The MRSG+erythromycin plates were then replica plated onto MRS with the sugar being assayed (cellobios, galactose, mannitol, mannose, or xylose) and onto MRSG. After overnight incubation, growth was compared on the selective plate and the glucose plate. Plates were imaged using the macroscope. Colonies that grew on MRSG but not on the selective plate were considered knockouts for the targeted sugar metabolism gene.

### Library Construction and Next Generation Sequencing

For sequencing, all the colonies from the initial selection plate (for *E. coli*) or the final MSRG replica plate (for *W. confusa*) were scraped off of the plate and re-suspended using 1 mL of ddH_2_O into 1.5 mL micro-centrifuge tubes. The cells were spun down and the supernatant was discarded. Pellet were stored at −20°C. Genomic DNA was extracted using Promega Genomic DNA Kit. *W. confusa* genomic DNA can be extracted using lysozyme, as per the Promega protocol for Gram-positive bacteria. Once extracted, the DNA was normalized to 10 ng*/*µL and used as a template for a PCR of the appropriate genomic region (see Supp. File A.3 for amplified regions). PCRs were purified using Zymo Clean and Concentrate and were normalized to 50 ng*/*µL. Sets of PCRs from different loci (e.g. the first replicates from all of the sugar genes in both *E. coli* and *W. confusa*) were combined. These were sheered on a M220 Focused-ultrasonicator (Covaris) with a target size of 500 bp. Size distributions were verified on a Bioanalyzer. The final concentration of DNA was generally low (e.g. 20 ng*/*µL).

A NEBNext Ultra II DNA Library Prep Kit was used for preparation of sheered DNA for sequencing. The initial amplification step was done with 20 cycles due to the low concentration of sheered DNA. Loss of diversity due to the large number of amplification cycles was not a concern due to the library sizes being small. Standard index primers, purchased from NEB, were used.

Samples were run on a MISeq. Before loading, library concentration was measured using qPCR, nanodrop, and qBIT. These gave varying concentrations, so the qPCR concentration was used for loading. Read density indicated the concentration was 1/4 of the concentration indicated by the qPCR (the nanodrop was, in fact, the most accurate).

Reads were aligned using Geneious software. Each replicate was aligned to its reference sequence allowing for discovery of any size deletions. Supplemental File A.4 contains a list of observed deletions supported by at least five reads for each gRNA and replicate.

### Confirmation of *W. confusa* knockout phenotypes

For confirmation of knockout genotype and sugar metabolism deficiencies in cellobios and mannose, *W. confusa* was re-transformed with the relevant plasmids. For each transformation, 16 colonies were re-struck onto MSRG+erythromycin plates. One colony from each re-streak was propagated and glycerol stocks were made. At the same time, 100 µL of each culture was spun down, decanted, and stored at −80°C.

For colony PCRs, frozen cell pellets were first re-suspended in 100 µL of pH 8.0 TE. 1 µL of each re-suspensions was used as template in 25 µL PCR reactions. Standard PCR protocol was used, except the initial 98°C denaturation step was extended to 5 minutes. Deletions of cellobios and mannose genes were confirmed by “primer-walking” from the PCR primers until the deletions were sequenced.

Sugar metabolism was confirmed by using the minimal media plates described above.

### Equipment

OD_600_ was measured using an Ultrospec 10 (Amersham Biosciences) and plastic cuvettes. Biorad thermocyclers were used, as were Eppendorf 5810 and 5810 R centrifuges. HT Multitron and Shell Lab Low Temperature Incubator were used for shaking and stationary incubation, respectively. A custom built macroscope, courtesy of the Kishony Lab and the Harvard Department of Systems biology (see: https://openwetware.org/wiki/Macroscope), was used to take pictures of agar plates. For plate reader experiments, a Biotek HT1 Synergy was used.

## Supporting information

Plasmid Sequences

## Acknowledgments

This work was performed during my time in, and funded by, Pamela A. Silver’s Laboratory at Harvard Medical School. T. W. Giessen, S. G. Hays, and B. B. Hsu provided thoughtful comments on this manuscript.

## Footnotes

† It is important to use a *temperature-controlled centrifuge* for these steps, as a “room temperature” centrifuge will become too hot and cause reduced cell viability.

