## Supplemental Tables and Figures for "Heterologous Cas9 and non-homologous end joining as a Potentially Organism-Agnostic Knockout (POAK) system in bacteria"

### Supplemental Figures and Information

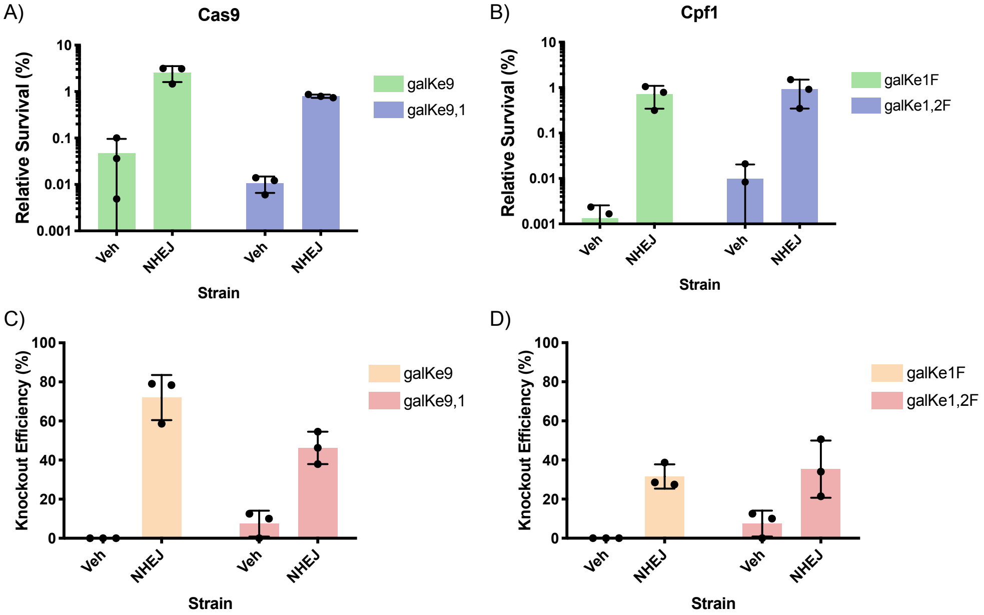


Figure 9: **Functioning of a dual plasmid Cas9/Cpf1 and NHEJ system in *E. coli*** **A)** Relative survival when *E. coli*, either with a blank vehicle plasmid (Veh) or an NHEJ expressing plasmid (NHEJ), is transformed with Cas9 with a single gRNA targeting *galK* (galKe9) or two gRNAs (galKe9,1). **B)** Relative survival when *E. coli*, either with a blank plasmid (Veh) or an NHEJ expressing plasmid (NHEJ), is transformed with Cpf1 with a single gRNA targeting *galK* (galKe1F) or two gRNAs targeting (galKe1,2F). **C)** Knockout efficiency from **A**. **D)** Knockout efficiency from **B**.

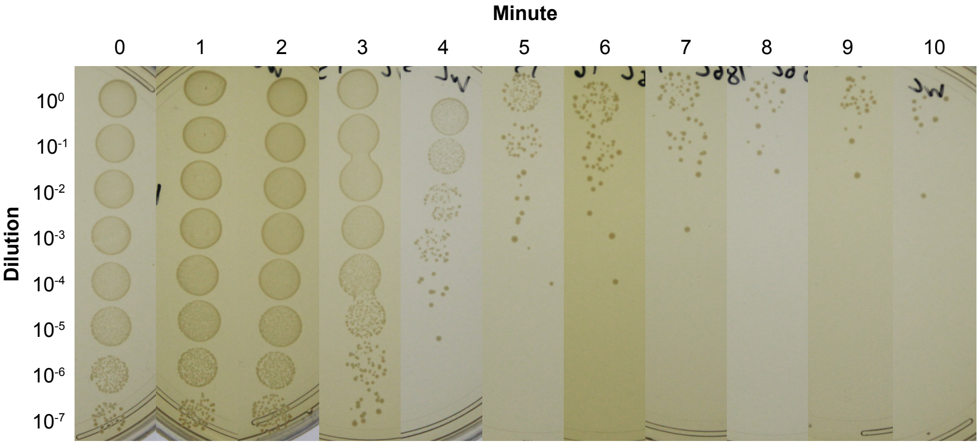


Figure 10: ***W. confusa* dies rapidly at** 56 °C A *W. confusa* culture was incubated at 56 °C for the number of minutes indicated, then serially diluted and spotted on MRS.

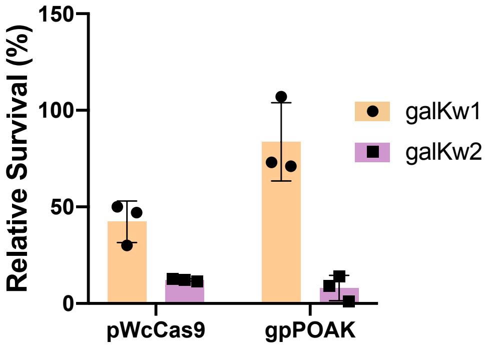


Figure 11: **Cas9 mediated death is gRNA dependent in *W. confusa*** Relative survival of *W. confusa* after transformation with pWcCas9 and gpPOAK with either the galKw1 gRNA or the galKw2 gRNA. The bar is the mean of three transformations, and error bars are standard deviation.

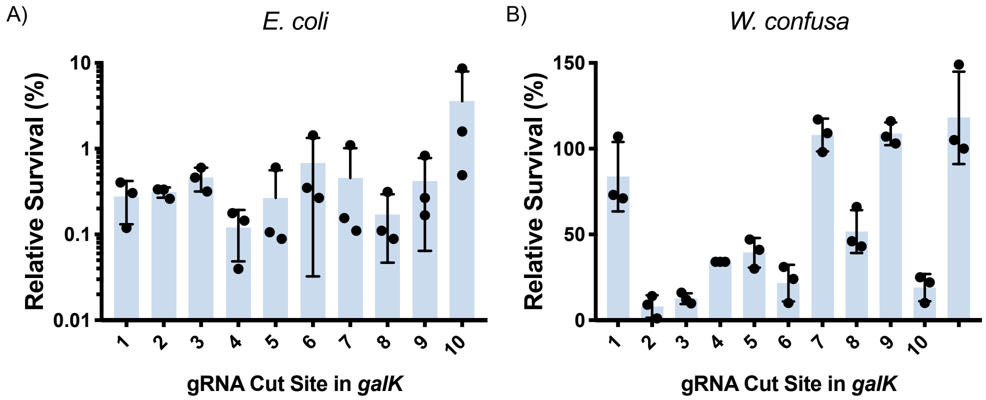


Figure 12: **Effect of cut position on relative survival in POAK knockouts in *E. coli* and *W. confusa*** **A)** Relative survival when *E. coli* is transformed with gnPOAK containing a gRNA that cuts at the indicated position of galK. Bars are the mean of three transformations, error bars represent the standard deviation. **B)** Relative survival when *W. confusa* is transformed with gpPOAK containing a gRNA that cuts at the indicated position of galK. Cut sites 115 bp and 213 bp are gRNAs galk1 and galk2 respectively. Bars are the mean of three transformations, error bars represent the standard deviation.

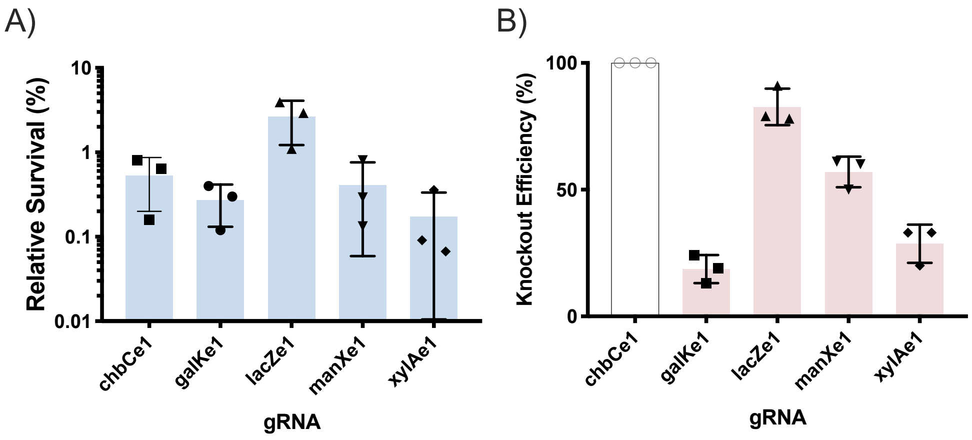


Figure 13: **Survival and knockout efficiency of gnPOAK for five *E. coli* genes A)** Relative survival when *E. coli* is transformed with gnPOAK containing gRNAs targeting 5 different genes. Bars are mean of three transformations, error bars are standard deviation. **B)** Knockout efficiency for the same five gRNAs. *chbC* knockouts could not be assayed, as the gene does not confer a measurable phenotype in MG1655[38]. Bars are mean of three transformations, error bars are standard deviation.

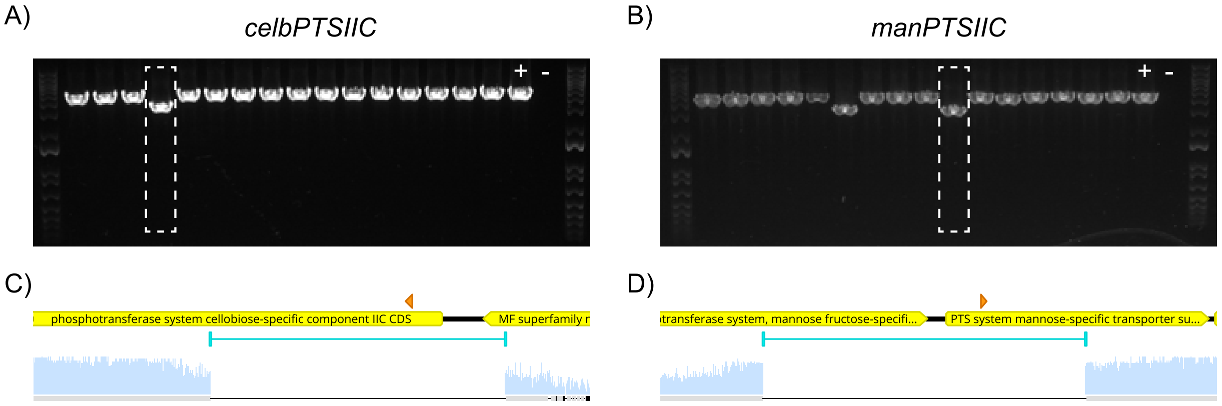


Figure 14: **Knockouts of *celbPTSIIC* and *manPTSIIC* in *W. confusa* A)** and **B)** PCRs to isolate knockouts in **A)** *celbPTSIIC* and **B)** *manPTSIIC* from 16 colonies transformed with the celbPTSIICw1 gRNA and the manPTSIICw1 gRNA respectively. Positive and negative template controls are shown in the last lanes. **C)** and **D)** Sequencing of highlighted PCRS from **A** and **B** respectively. Orange arrows represent the cut sites, and the regions with no light blue sequencing coverage delineate the deleted sequences.

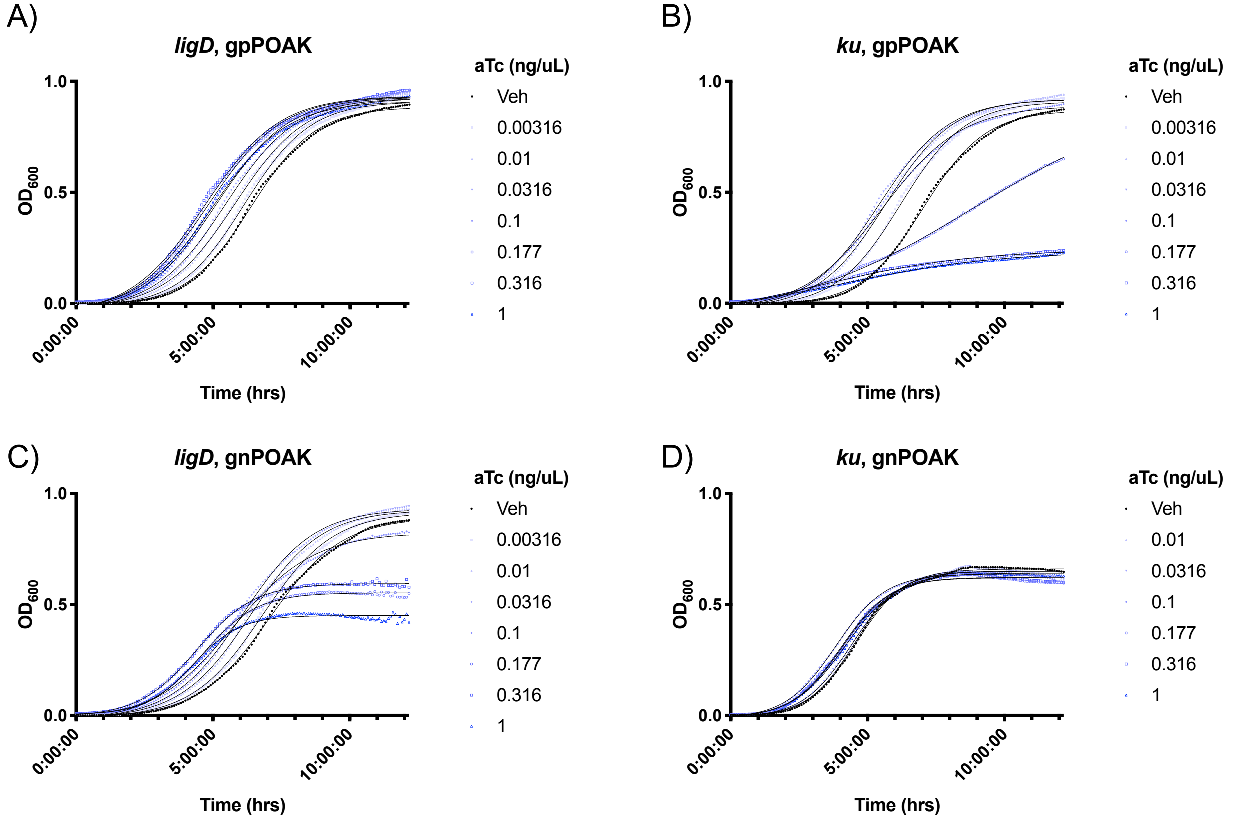


Figure 15: **Ku expression reduces growth in *E. coli*** *E. coli* was grown with either a gpPOAK or gnPOAK plasmid that had an incomplete NHEJ system, such that one gene was deleted and the other (*ku* or *ligD*) was under $P_{tetA}$ regulation. The *E. coli* was grown with selection as well as different concentrations of inducer (aTc). **A)** gpPOAK with only *ligD*. **B)** gpPOAK with only *ku*. **C)** gnPOAK with only *ligD*. **D)** gnPOAK with only *ku*.

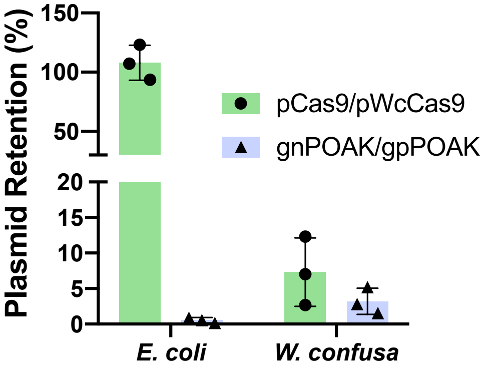


Figure 16: **gnPOAK is unstable in *E. coli* and both pWcCas9 and gpPOAK are unstable in *W. confusa*** *E. coli* and *W. confusa* were grown with aTc and the relevant plasmid (pCas9 and gnPOAK for *E. coli*, pWcCas9 and gpPOAK for *W. confusa*). Cultures were plated on selective (for plasmid) and non-selective plates, and the percentage of colonies that grew on the selective places relative to non-selective is reported. Bars are the mean of three different cultures, and error bars represent the standard deviation.

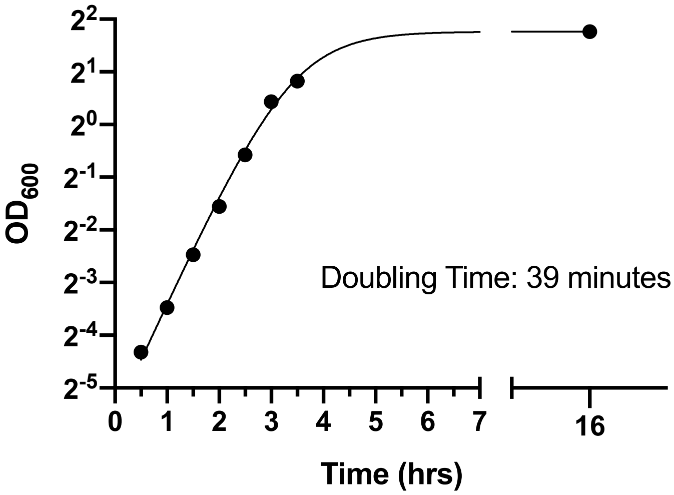


Figure 17: ***W. confusa* doubles every 40 minutes** Representative growth curve of *W. confusa* when grown in MRSG at 37 °C. A four-parameter logistics curve is shown and was fit using Prism.

### Strains, Plasmids, and gRNAs

Table 4: **Strains**

| Strain | Organism | Genotype |
|---|---|---|
| MG1655 | *Escherichia coli* | Wild Type |
| DSM 20196 | *Weissella Confusa* | Wild Type |
| BW25113 | *Escherichia coli* | $\Delta$(*araD-araB*)567, $\Delta$*lacZ*4787 (::rrnB-3), $\lambda$-, rph-1, $\Delta$(*rhaD-rhaB*)568, *hsdR*514 |
| JW1726-1$\Delta$*kanR* | *Escherichia coli* | BW25113 $\Delta$*chbC*513 |
| JW0740-3$\Delta$*kanR* | *Escherichia coli* | BW25113 $\Delta$*galk*729 |
| JW1806-1$\Delta$*kanR* | *Escherichia coli* | BW25113 $\Delta$*manX*741 |
| JW3537-1$\Delta$*kanR* | *Escherichia coli* | BW25113 $\Delta$*xylA*748 |

Table 5: **Plasmids**

| Plasmid | Description | Strain | Resistance | Host |
|---|---|---|---|---|
| pVeh | Empty vector (i.e. Veh) | IP_8 | Kn | Turbo |
| pNHEJ | pVeh+Ptrc(NHEJ) | IP_22 | Kn | Turbo |
| pCas9sp | PproC(Cas9) | IP_23 | Sp | Δgalk |
| pCas9spgalK9,1 | PproC(Cas9), galk9,1 | IP_24 | Sp | Δgalk |
| pCas9spgalK9 | PproC(Cas9), galk9 | IP_25 | Sp | Δgalk |
| pCpf1sp | PproC(Cpf1) | IP_34 | Sp | Δgalk |
| pCpf1spgalK1F | PproC(Cpf1), galk1F | IP_36 | Sp | Δgalk |
| pCpf1spgalK1,2F | PproC(Cpf1), galk1,2F | IP_37 | Sp | Δgalk |
| gpPOAKtemp | PproC(Cas9), Pteta(ligd, ku), pBAV1K backbone | IP_104 | Erm | DH5$\alpha$ |
| pCas9temp | PproC(Cas9), pBAV1K backbone | IP_109 | Erm | DH10ß |
| pWcCas9 | Pwc-eno(Cas9), pBAV1K backbone | IP_128 | Erm | DH10ß |
| gpPOAK | Pwc-eno(Cas9), Pteta(ligd, ku), pBAV1K backbone | IP_132 | Erm | DH10ß |
| pCas9 | PproC(Cas9), pBBR1 origin, KnR | IP_245 | Kn | ΔgalK |
| gnPOAK | PproC(Cas9), Pteta(ligd, ku), pBBR1 origin, KnR | IP_246 | Kn | ΔgalK |
| pCpf1 | PproC(Cpf1), pBBR1 origin, KnR | IP_247 | Kn | ΔgalK |
| gnPOAK_Cpf1 | PproC(Cpf1), Pteta(ligd, ku), pBBR1 origin, KnR | IP_248 | Kn | Δgalk |
| pCas9galK9 | | IP_250 | Kn | ΔgalK |
| pCas9galK9,1 | | IP_251 | Kn | Δgalk |
| gnPOAKgalK9 | | IP_252 | Kn | ΔgalK |
| gnPOAKgalK9,1 | | IP_253 | Kn | Δgalk |
| pCpf1galK1F | | IP_255 | Kn | ΔgalK |

Table 5: (continued)

| Plasmid | Description | Strain | Resistance | Host |
|---|---|---|---|---|
| pCpf1galK1,2F | | IP_256 | Kn | ∆galk |
| gnPOAK_Cpf1galK1F | | IP_259 | Kn | ∆galK |
| gnPOAK_Cpf1galK1,2F | | IP_260 | Kn | ∆galk |
| gpPOAKcelbPTSIICw1 | | IP_281 | Erm | DH10ß |
| gpPOAKgalKw1 | | IP_282 | Erm | DH10ß |
| gpPOAKgalKw2 | | IP_283 | Erm | DH10ß |
| gpPOAKgalKw3 | | IP_284 | Erm | DH10ß |
| gpPOAKgalKw4 | | IP_285 | Erm | DH10ß |
| gpPOAKgalKw5 | | IP_286 | Erm | DH10ß |
| gpPOAKgalKw6 | | IP_287 | Erm | DH10ß |
| gpPOAKgalKw7 | | IP_288 | Erm | DH10ß |
| gpPOAKgalKw8 | | IP_289 | Erm | DH10ß |
| gpPOAKgalKw9 | | IP_290 | Erm | DH10ß |
| gpPOAKgalKw10 | | IP_291 | Erm | DH10ß |
| gpPOAKgalKw11 | | IP_292 | Erm | DH10ß |
| gpPOAKmaltPw1 | | IP_293 | Erm | DH10ß |
| gpPOAKmanPTSIICw1 | | IP_294 | Erm | DH10ß |
| gpPOAKxylAw1 | | IP_295 | Erm | DH10ß |
| gnPOAKchbCe1 | | IP_296 | Kn | ∆celB |
| gnPOAKgalKe1 | | IP_297 | Kn | ∆galk |
| gnPOAKgalKe2 | | IP_298 | Kn | ∆galk |
| gnPOAKgalKe3 | | IP_299 | Kn | ∆galk |
| gnPOAKgalKe4 | | IP_300 | Kn | ∆galk |
| gnPOAKgalKe5 | | IP_301 | Kn | ∆galk |
| gnPOAKgalKe6 | | IP_302 | Kn | ∆galk |
| gnPOAKgalKe7 | | IP_303 | Kn | ∆galk |
| gnPOAKgalKe8 | | IP_304 | Kn | ∆galk |

Table 5: (continued)

| Plasmid | Description | Strain | Resistance | Host |
|---|---|---|---|---|
| gnPOAKgalKe9 | | IP_305 | Kn | Δgalk |
| gnPOAKgalKe10 | | IP_306 | Kn | Δgalk |
| gnPOAKlacZe1 | | IP_307 | Kn | Δgalk |
| gnPOAKmanXe1 | | IP_308 | Kn | ΔmanY |
| gnPOAKxylAe1 | | IP_309 | Kn | ΔxylA |
| pWcCas9galKw1 | | IP_315 | Erm | DH10ß |
| pCas9galKe1 | | IP_317 | Kn | Δgalk |
| pWcCas9galKw2 | | IP_336 | Erm | DH10ß |

Table 6: **gRNAs**

| Name | Sequence | Cut Site from ATG (bp) |
|---|---|---|
| galK1F | CCAACGCATTTGGCTACCCTGC | 34 |
| galK2F | TAAACCATCACAAGGAGCAGGA | 1118 |
| celbPTSIICw1 | AATTGCGTATGCGATGCCAT | 112 |
| galKw1 | GAGCACACCGATTATAATGG | 115 |
| galKw2 | CTATTCAGCGAACTTCCCAG | 213 |
| galKw3 | TTGGCTACCCCGTAACAAAA | 332 |
| galKw4 | AATATGCTAGCTGACTTGAT | 433 |
| galKw5 | GGCCTGTTCGTCTTCACCAA | 557 |
| galKw6 | AAAGTACAACGAACGTCGTG | 684 |
| galKw7 | GGCAGCGTCAGCGTCATTAA | 786 |
| galKw8 | GCCCAAACGTTTCCAAATCG | 897 |
| galKw9 | GGTGCTCGTATGACTGGTGC | 1021 |
| galKw10 | ATTTCGGCGACGAAAAATGA | 1144 |
| galKw11 | AGATTCACATAGTAGTCAAT | 1208 |
| maltPw1 | AATGAGTACATGGGTATGCG | 116 |
| manPTSIICw1 | ATTGGCAACTGGTCACCTAA | 127 |
| xylAw1 | CCTAAGGTAGAATTTATCGG | 119 |
| chbCe1 | AATGCCGTTAACCCTTGCGG | 124 |
| galKe1 | GTTCACCAATCAAATTCACG | 93 |
| galKe2 | CTGCCATCACGCGAACTTTA | 195 |
| galKe3 | GGCTAACTACGTTCGTGGCG | 285 |
| galKe4 | GCTTCACTGGAAGTCGCGGT | 400 |
| galKe5 | GATCAGCTAATTTCCGCGCT | 529 |
| galKe6 | CAGTAACTTCAAACGTACCC | 642 |
| galKe7 | AACAGCGTTGAACTCTTCAA | 749 |
| galKe8 | GCAAGGCGACCTGAAACGTA | 858 |
| galKe9 | GGTGGCGTACGCATGACCGG | 988 |

Table 6: (continued)

| Name | Sequence | Cut Site from ATG (bp) |
|---|---|---|
| galKe10 | GAACAATATGAAGCAAAAAC | 1081 |
| lacZe1 | TTACGCCAGCTGGCGAAAGG | 96 |
| manXe1 | GTACTTTTCAATCAGCGTTT | 127 |
| xylAe1 | CTTACCCAACACCAGTTCGT | 88 |

### gRNA Design and Cloning (for spCas9)

1. Use a computer to design gRNAs. PAM: 5' NGG. The length of the gRNAs doesn't really matter. I have used 22 bp before, but now usually use 20 bp because I arbitrarily decided to.

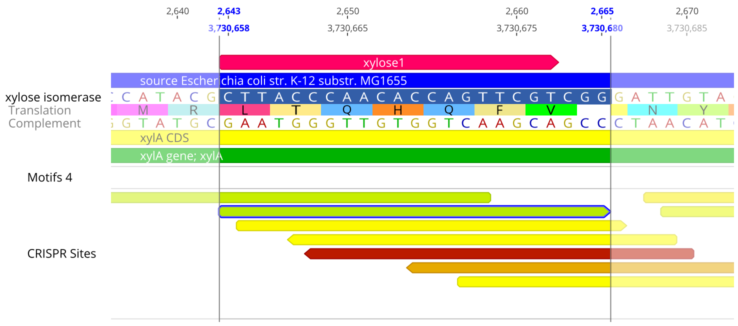


Figure 18: gRNA of interest (in green)

2. Select the targeting region of the gRNA (everything besides the NGG)

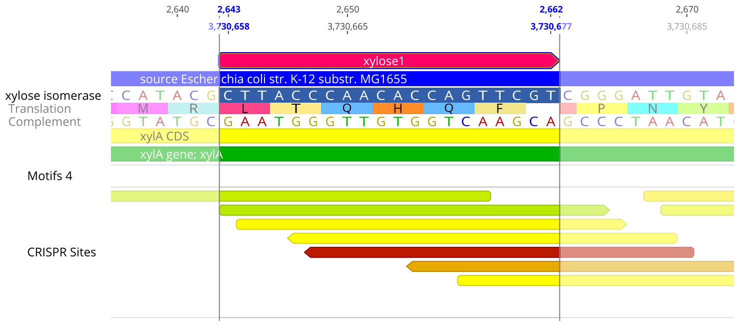


Figure 19: Portion of desired gRNA to be copied (in pink)

3. Copy it into the gRNA template such that the 5' side of the gRNA is followed by a GTTT.

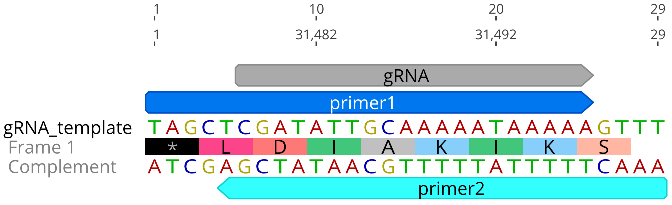


Figure 20: gRNA template

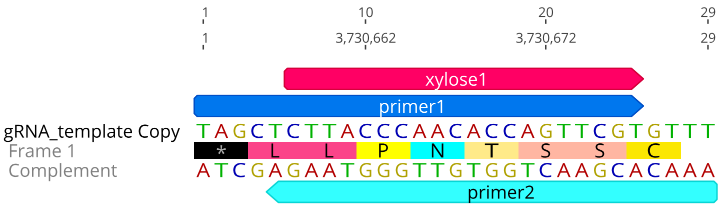


Figure 21: Example gRNA copied into template

4. Order the oligos labeled as "primer 1" and "primer 2"

5. A day or two before you are going to be doing the cloning, make the following reaction:

Table 7: Reaction for Oligo Phosphorylation and Annealing

| | |
|---|---|
| 1 µL | T4 Polynucleotide Kinase |
| 2 µL | 100 µM Oligo 1 |
| 2 µL | 100 µM Oligo 2 |
| 4 µL | 10x T4 Ligase Buffer |
| 31 µL | dd$H_2O$ |

6. And use the following conditions

Table 8: Thermocycle for Oligo Phosphorylation and Annealing

| | |
|---|---|
| 37 °C | 30 min. |
| 95 °C | 5 min. |
| −1 °C/min | |
| 25 °C | 1 min. |
| 4 °C | ∞ |

7. Dilute reaction 1:50 in water, and use 1 µL of dilution per 20 µL golden gate reaction.
